# Citizen science data enhance spatio-temporal extent and resolution of animal population studies

**DOI:** 10.1101/352708

**Authors:** Catherine C. Sun, Angela K. Fuller, Jeremy E. Hurst

## Abstract

Informed management and conservation decisions for animal populations often require data at sufficient geographic, temporal, and demographic resolutions for precise and unbiased estimates of parameters including population size and demographic rates. Recently developed integrated population models estimate such parameters by unifying population presence-absence and demographic data, and we demonstrate how citizen science offers a cost-efficient mechanism to collect such data. We describe the early results of iSeeMammals, a citizen science project that collects opportunistic data on the black bear population in New York State by enlisting volunteers to collect data through observations, hikes, and trail cameras. In 10 months, iSeeMammals increased the spatio-temporal extent of data collection by approximately fourfold and reduced cost by 83% compared to systematic sampling. In combination with other datasets in integrated population model frameworks, large, spatiotemporally extensive datasets from citizen science projects like iSeeMammals can help improve inferences about population-level structure and dynamics.

## Introduction

Successful conservation and management of animal populations requires knowledge about population abundance, distribution, and dynamics such as growth (Nichols and Williams, 2006; Williams et al., 2002). These are critical for assessing population status and predicting population viability and changes (Beissinger and Westphal, 1998). This is especially important as species extinction risk is tied to the loss of discrete populations, with recent studies demonstrating range contractions of 94-99% for some large carnivores (Wolf and Ripple, 2017). Accelerating declines in animal populations worldwide necessitate methods to reliably and efficiently estimate population parameters such as population size, and vital rates such as survival probability and recruitment rate. Data on individuals at high spatial and temporal resolutions from across species’ ranges are needed to detect patterns across landscapes (Ashcroft et al., 2009), but limited resources often lead to datasets that cover an area too small to represent the species’ range or are too sparse to provide precise and accurate estimates of population parameters. As a result, unfortunately, many species of conservation concern are managed without sufficient information on population status or trends.

Precise estimates of population parameters are necessary for conservation decisions to discriminate between populations that do and do not require conservation action. Spatial capture-recapture (SCR) approaches estimate population size and structure by tabulating the times and locations at which individuals are detected and accounting for detection probability less than 100%. When collected over timeframes in which individuals are born or die, i.e., a robust design (Pollock, 1982), the data provide information about vital rates such as survival and recruitment. However, the intense sampling effort required to repeatedly detect individuals limits the spatiotemporal extent of SCR approaches and inferences about population demographics. While datasets that are low resolution or spatiotemporally limited may not by themselves contain enough information to accurately and precisely identify and predict landscape-wide population patterns, they can be merged with other datasets of greater resolution or coverage (i.e., different scales) in integrated population models (IPMs). IPMs model population structure and demographics by uniting multiple datasets, making it possible to estimate parameters not previously identifiable by the individual datasets and achieve more precise and unbiased estimates of abundance and population dynamics (Chandler and Clark, 2014). A hierarchical framework based on SCR data enables direct inference on population parameters, by modeling individuals on the landscape as a spatial point process, using a detection sub-model to account for imperfect detection and individual heterogeneity (Royle and Young, 2008). Integrating species-level data, which contain less direct information about population demographics but can cover large spatiotemporal extents, can therefore improve, expand, and scale up population inferences (Chandler and Clark, 2014).

Citizen science, in which members of the public participate in scientific research (Bonney et al., 2009; Dickinson et al., 2012; McKinley et al., 2017), is a potential source of such spatiotemporally extensive data. With the aid of technology, citizen science has been used to study a variety of ecological processes, including species distribution, phenology, and behavior (Dickinson et al., 2012). Citizen science approaches commonly collect data on the repeated presence and absence (P-A data) of species at various locations as well information only on the presence (P-0 data) of species at single points in time (Pocock et al., 2017). Furthermore, citizen science data collected opportunistically, or at un-predetermined times and locations, which may include point counts (Wood et al., 2011), species checklists (Kéry et al., 2010), and SCR data (Tenan et al., 2017), can produce robust inferences equivalent to those derived from traditional, systematic sampling at predetermined times and locations (Nagy et al., 2012). Large citizen science data sets collected at large spatiotemporal extents may also reduce the amount of un-sampled population and landscape variation, lowering the risk of false inferences (Bain et al., 2015) that can affect extrapolation with scale-limited datasets and lead to misguided conservation actions. Integrating citizen science data into IPMs may therefore yield precise, and unbiased estimates of abundance, distribution, and vital rates of wildlife populations of conservation concern. The number of published articles regarding citizen science and its use have increased since the 1990s (Follett and Strezov, 2015). Yet there are relatively few examples of its application in IPMs. To illustrate how species-level data can be collected with citizen science approaches for use in IPMs to improve population inferences for species conservation and management, we describe the development and application of a citizen science project called iSeeMammals. iSeeMammals enlists volunteers to submit opportunistic data on black bears *(Ursus americanus)* to study their populations in New York State, and supplements ongoing systematic SCR data collection. Previous studies in New York and elsewhere have been conducted only at small spatial extents (Belant et al., 2005; Drewry et al., 2013; Sun et al., 2017). We discuss how large and spatiotemporally extensive datasets from citizen science projects like iSeeMammals could be applied in IPMs to estimate population parameters and trends relevant to management and conservation.

## Materials and Methods

iSeeMammals launched in 2017, and allows users to opportunistically collect spatially-referenced data with one-time observations, hikes, and trail cameras. To collect data, we developed a website (iseemammals.org) and smartphone application (app) that is available for free download (Figure 1). iSeeMammals seeks members of the public who hike, hunt, track, and partake in general outdoor, wildlife recreational activities.

**Figure 1.**
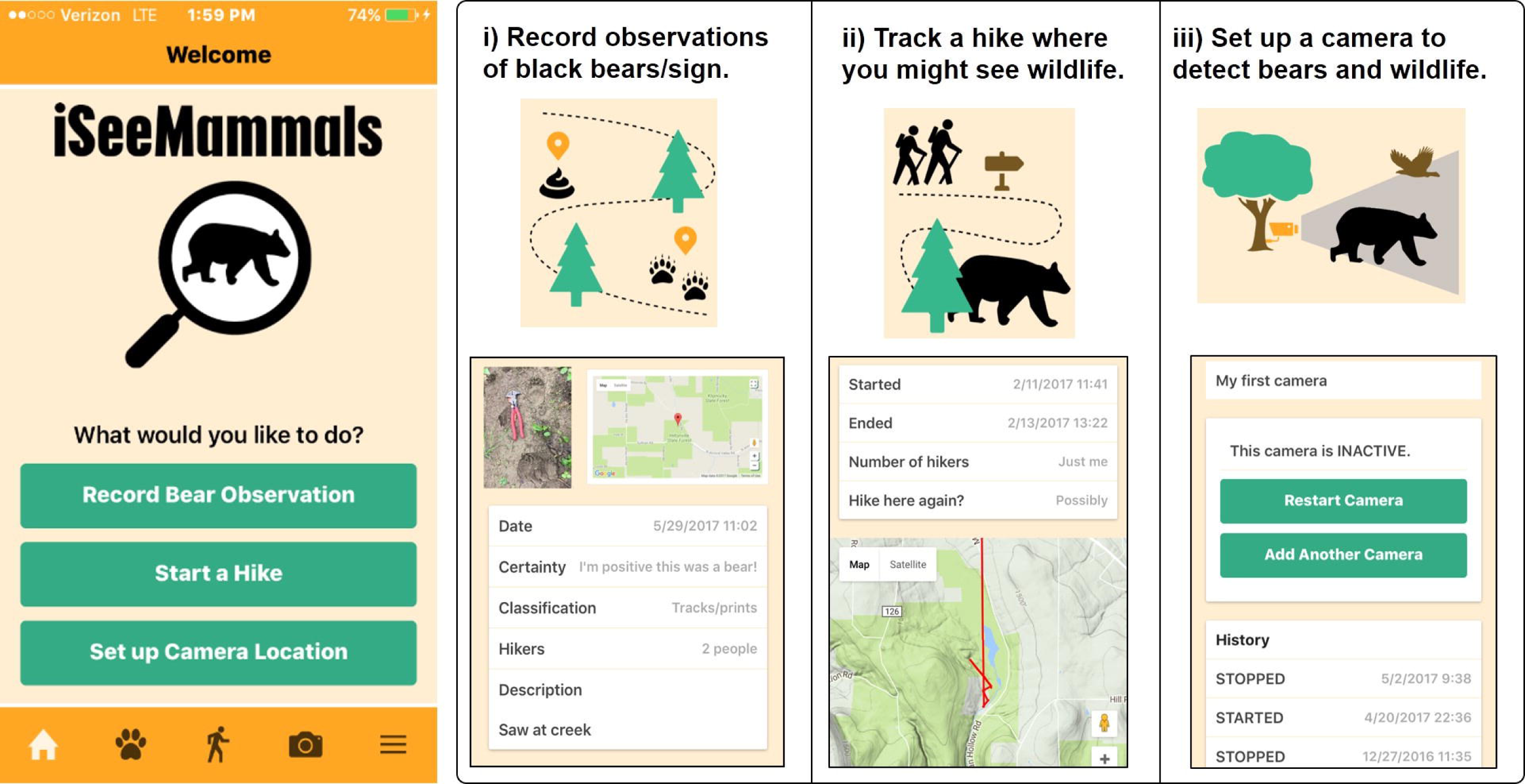
Home screen of iSeeMammals smartphone app and the three features for collecting presence-only (P-O) data of bears or bear signs, or presence-absence (P-A) data with observations, hikes, and trail cameras.

One-time observations of bear or bear sign (scat, tracks, hair, markings) provide presence-only (P-O) data. Users may include a photograph, and then confirm the time and GPS location of the observation, which may be obtained from the photograph metadata. A text description and questions about the observation type (bear, scat, track, hair, or marking), confidence in identification, and number of people present for the observation supply information for filtering data and quantifying detection effort.

Hikes and cameras provide presence-absence (P-A) data by explicitly including where and when no observations are recorded. Users collect GPS hike coordinates with the app at approximately 500 m increments to prevent excessive battery drainage during long hikes. Users attach any observations taken during the hike, confirm the general accuracy of the route, and provide information about the number of hikers. Users also report the likelihood of returning at a later date to repeat the hike, which would provide temporal replication and therefore improve estimates of detection probability and population parameters.

From trail cameras, users collect the cameras’ GPS coordinates with the app and maintain a history log of camera activity. Additional information is collected about the camera brand, model, times of day the camera was operational, periods of camera malfunction, and whether or not bears were detected on the camera. Users are instructed to submit bear photographs from the camera through the website. Absence data from cameras were inferred from lack of bear photographs.

We summarized the iSeeMammals data collected between January 1 – October 31, 2017. We rejected one-time observations lacking spatial data or if species identification was incorrect based on the provided photograph, and removed duplicates based on photographs and descriptions. We rejected hikes that were either described as inaccurate by the user, were less than one minute, or had only 1 set of GPS coordinates. We rejected camera data if they monitored a location for less than 1 day. We “ended” ongoing camera on October 31, 2017 and assumed no malfunctions and no pictures of bears were collected. For each collection method, we calculated summary statistics and describe data filtering, data quality, and spatial patterns.

## Results

A total of 712 users registered with iSeeMammals in 10 months. Of those, 624 users (88%) activated their accounts, which involved clicking on an email link sent after registration. At least 126 users (18%) submitted a total of 471 independent sets of spatial data in New York, with the majority being one-time observations (79%).

Citizen scientists submitted 373 one-time observations (Table 1). All types of bear sign were reported. We rejected 14 misidentifications, 8 duplicates, 10 with no spatial information, and 2 that were rescinded by citizen scientists due to incorrect information. A significantly greater proportion of accepted observations reported confident identification (1.0) compared to rejected observations (0.72) (Fisher’s exact test: 2-tailed p<0.001). Across 38 counties in New York and within a minimum convex polygon (MCP) of 113,392 km^2^, iSeeMammals received 290 observations (86%). All observations reported confident identification and 222 observations (77%) included photographs, although 25 photographs submitted as observations were from trail cameras of unregistered users. A significantly greater proportion of observations with photographs reported bear signs rather than bears (0.41) compared to observations without photographs (0.06), (Fisher’s exact test: 2-tailed p<0.001). Most observations (88%) were submitted by parties of 1 or 2 people (150 and 105 observations, respectively) but parties of ≥5 people also submitted 10 observations (3.4%).

**Table 1.** iSeeMammals one-time observations, hikes, and trail camera data received during a 10 month period (January 1 and October 31, 2017)^1^.

| Data type | Data class | Total submissions | Accepted NY submissions |
| --- | --- | --- | --- |
| One-time observation (118) | Presence-only | 373 (9%<sup>®</sup>) | 290 |
| *Bear* |  | 221 | 194(130) |
| *Scat* |  | 108 | 63(59) |
| *Track* |  | 16 | 8(8) |
| *Hair* |  | 5 | 4(4) |
| *Markings* |  | 23 | 21(21) |
| Hike (4) | Presence-absence | 103 (56%) | 43 |
| *Bear* |  | 17 | 17 |
| *Scat* |  | 12 | 11 |
| *Track* |  | 0 | 0 |
| *Hair* |  | 0 | 0 |
| *Markings* |  | 3 | 3 |
| Trail camera (13) | Presence-absence | 81 (5%) | 63 (33) |
| Total (126) |  | 571 (70%) | 406 |

<sup>1</sup> Per data type, the number of users are shown in parentheses. The number of total submissions are shown, with the percentage filtered out shown in parentheses. For trail cameras, numbers are of locations at which data were collected. <sup>®</sup>Accepted and rejected observations had similar proportions of bear to bear signs (0.60 vs. 0.47, p-value=0.15). Accepted observations in New York are shown, for which the number that included photos are shown in parentheses.

Citizen scientists submitted 103 hikes, but we rejected 58 hikes (56%) (Table 1). Across 8 counties in New York and within an MCP of 25,400 km^2^, 43 hikes logged a total of 82.3 hours, with an average hike time of 1.9 hours (maximum 4.7 hours). Hikes collected an average of 24 GPS locations (range: 3-138) (Figure 2). The majority of hikes (37 hikes, 80%) had only 1 person, and most users indicated they would likely return to hike again in 3 months (38 users, 83%). A total of 18 hikes (42%) attached observations, with an overall average of 0.67 observations per hike. All hike observations were submitted with confidence. This resulted in 1,264 correlated spatial P-A data points.

**Figure 2.**
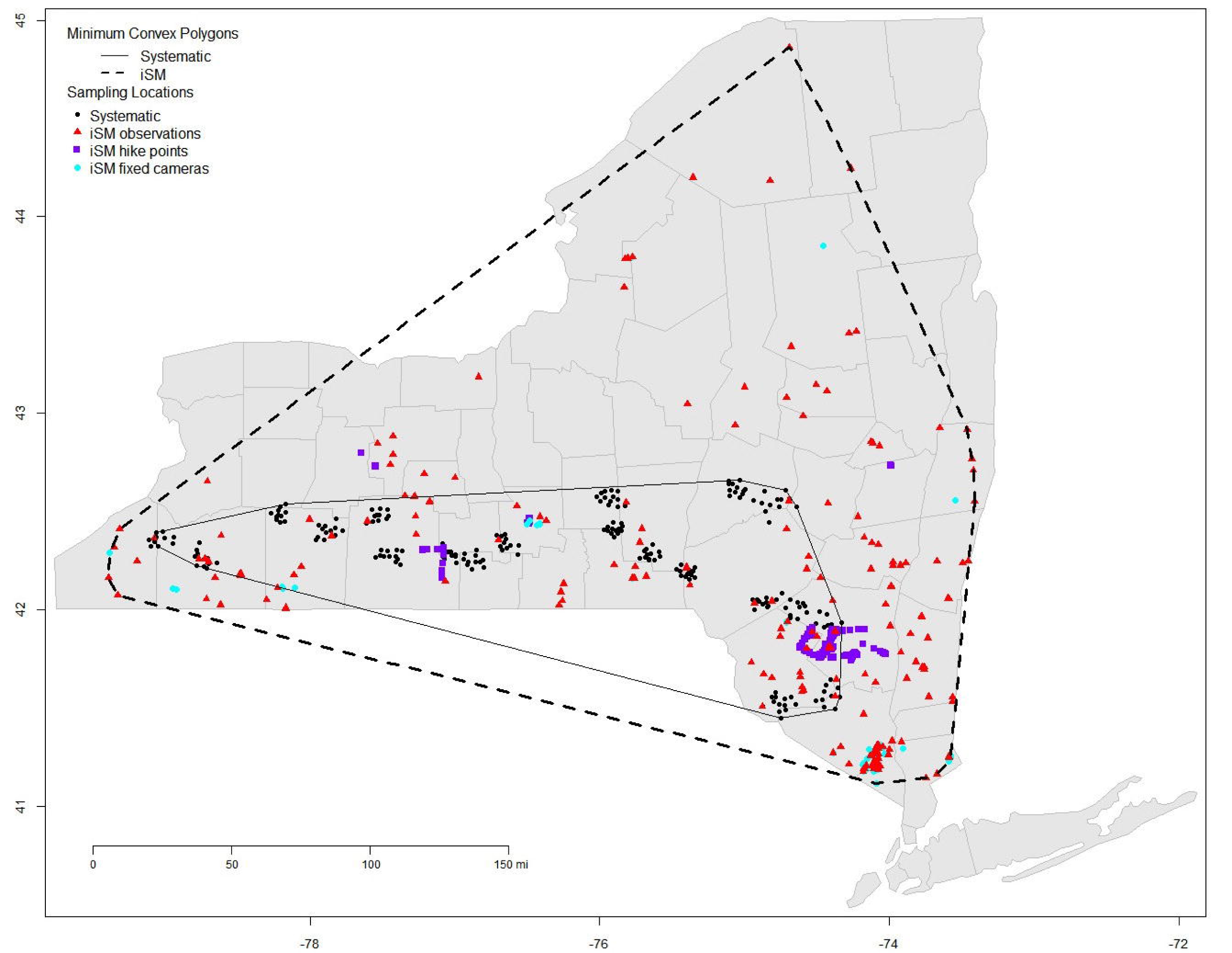
Location of iSeeMammals (iSM) one-time observations, hikes, and fixed cameras collected between January 1, 2017 and October 31, 2017. Also shown in black dots are the location of 241 systematic sampling sites using barbed wire collected in the summer of 2017. The minimum convex polygons for the systematic and iSeeMammals sampling, highlight the value of citizen science data for increasing spatial extent and quantity of data that can be collected. Coordinates shown in latitude and longitude.

Citizen scientists monitored 76 cameras at 81 different locations (Table 1). All cameras operated continuously. Across 12 counties in New York and within an MCP of 86,372 km^2^, 60 cameras were placed at 63 locations (Figure 2). Within New York, 2 cameras malfunctioned a total of 14 days, and 4 cameras were operational for less than 1 day (3%), resulting in 350 camera days collected. Cameras operated an average of 57 days per location (range: 6 - 153 days). iSeeMammals received 834 photographs of bears from 33 camera locations in New York, providing presence-absence information with an average of 25 photographs per location (range: 1 – 134 photographs), while the remaining 30 of 63 cameras did not detect black bears and therefore provided only absence data.

## Discussion

Citizen science can generate large quantities of spatiotemporally extensive, opportunistic data on the presence and absence of wildlife. Compared to systematic sampling conducted by researchers in 2017, iSeeMammals increased the quantity of data collection by 6.8-fold, the spatial extent by 3.7-fold (Figure 2), and the temporal extent by fourfoto January–October. Such data collection would not have been possible with limited systematic sampling. In the future, the proportional contribution of data by iSeeMammals may be even greater when systematic sampling is conducted less frequently or intensively and also as iSeeMammals becomes more widely used. iSeeMammals provides proof of concept that citizen science projects can be a useful alternative to biologist-driven data collection when collecting P-A data on species of conservation or management concern.

To use data collected in projects such as iSeeMammals as standalone datasets would be a narrow interpretation of what citizen science datasets can contribute to studying animal populations. Instead, incorporating citizen science datasets into integrated population models (IPMs) can improve estimates and inferences on population abundance, growth, and distribution, by covering larger regions to better represent the population’s spatial heterogeneity as well as longer time periods to detect more individuals and birth and death events (Figure 3). IPMs are an area of current and exciting development and there is increasing recognition of the value of incorporating citizen science data types into IPMs

**Figure 3.**
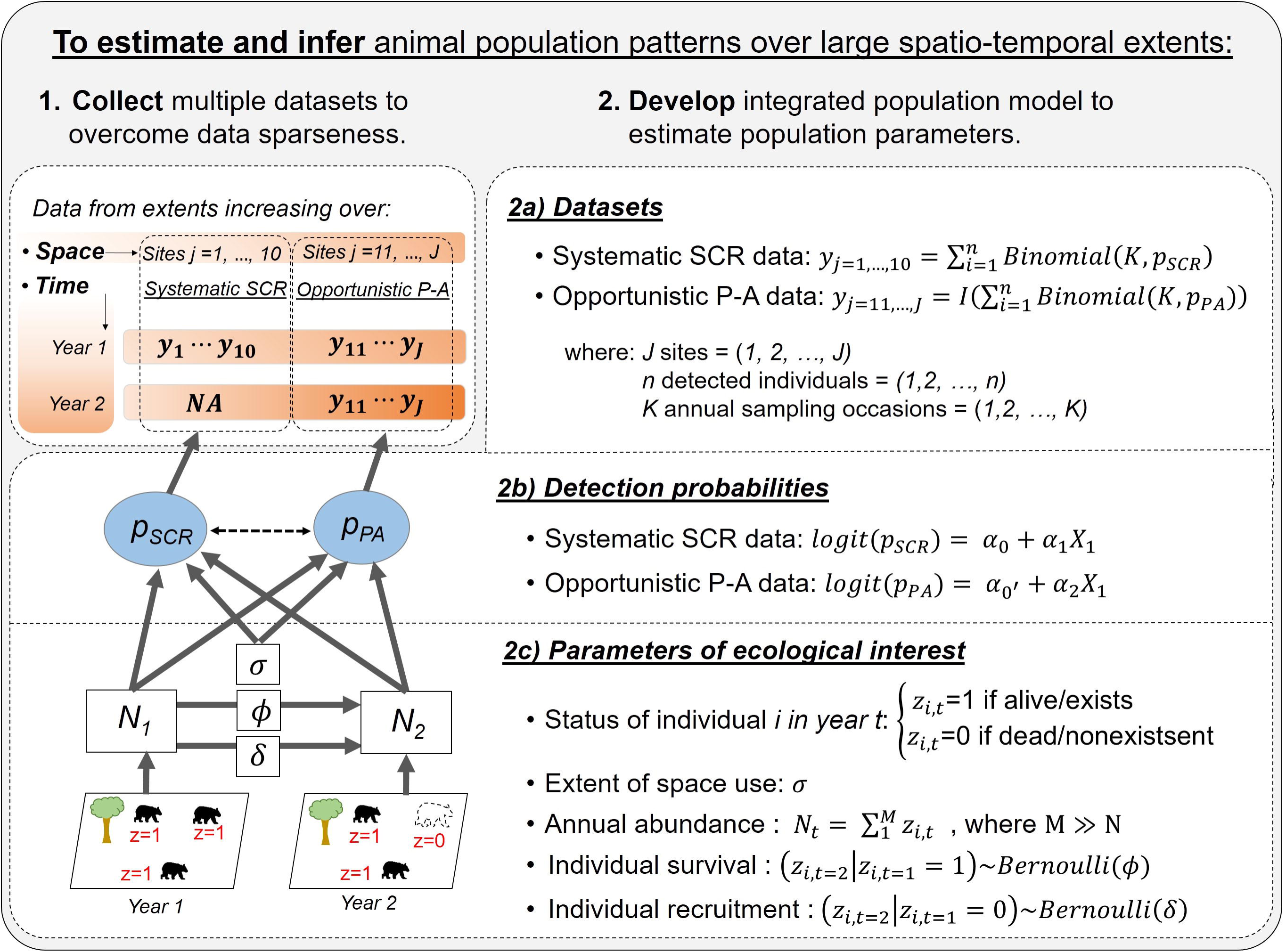
Framework for how collecting opportunistic data with citizen science approaches can contribute to the estimation of population parameters for informing animal conservation or management. Supplementing systematic data with opportunistic data, particularly presence-absence (PA) and presence-only (P-O) data, can expand the spatio-temporal extent of data collection and inference beyond what is possible with systematic data alone. Models that accommodate citizen science data are a current area of development (Sun et al. in review). A hierarchical formulation based on individuals, linked to the observed data through detection probabilities, will allow for estimation of abundance and demographic rates.

(Zipkin and Saunders, 2018). As iSeeMammals has demonstrated, P-A and P-0 datasets can be large and some citizen science platforms such as iNaturalist solely collect opportunistic P-0 data. Some IPMs have already incorporated opportunistic SCR data (Tenan et al., 2017), species-level count data (Robinson et al., 2018), and systematic species-level P-A data (Chandler and Clark, 2014), but the authors are unaware of any current IPMs that integrate opportunistic species-level P-A or P-0 data. It would be straightforward to develop a model with opportunistic P-A data that are collected independently of systematic SCR data (Sun et al. in review), in which the detection processes by which the opportunistic and systematic datasets are collected are either related in some fashion or entirely independent. Furthermore, formalizing the relationship between P-0 and P-A data in an IPM would open many citizen science P-0 datasets to robust population modeling rather than resort to approaches that generate pseudo-absences (Yackulic et al., 2013). An integrated modeling framework could also facilitate the joining of datasets from multiple citizen science platforms. Sensitivity analyses through simulations with the IPMs could also help determine the amounts of systematic and opportunistic data necessary to improve estimates or detect changes in population trends. By finding creative ways to use opportunistic datasets and developing IPMs that incorporate them, we may expand the reach of citizen science beyond exploratory investigations (Dickinson et al., 2010) to improve population estimates and prudently manage animal populations of conservation concern.

The process of filtering iSeeMammals data highlights the importance of maintaining citizen science data quality and quantity. To ensure data quality, approximately 67% of submitted data were filtered out. At the same time, losing absence data, which iSeeMammals explicitly collects, negatively impacts the estimation of detection probability and subsequently population parameters. Clearer guidelines for data collection, automatic reminder notifications, and improved GPS technologies on smartphones would help further improve the quality of spatial data. Continued and strategic outreach and communication

are also necessary to recruit new citizen scientists, maintain engagement, and target participation where data gaps exist. Only 18% of registered users (126 of 712) submitted iSeeMammals data, underscoring the need to understand the motivations for and against participation at all stages of engagement (Beirne and Lambin, 2013; Eveleigh et al., 2014; Nov et al., 2014). Incentives and the transformation of data collection tasks into games have proven to help retain participation and increase data quantity ((Bowser et al., 2013; Iacovides et al., 2013; Xue et al., 2016)).

Citizen science projects such as iSeeMammals may yield sampling and financial efficiencies that make it a cost-efficient approach to studying and monitoring population-level patterns. iSeeMammals received data from other northeastern states and on non-target wildlife species, suggesting initiatives could formally collect data across even larger extents and on more species. Furthermore, the development of iSeeMammals cost $32,000 in contrast to the annual cost of approximately $191,000 for systematic sampling, which includes genetic analyses, equipment, salary, and travel costs for approximately a dozen researchers and technicians. Once the infrastructure for a citizen science program has been established, opportunistic data collection may continue when and where systematic sampling is not possible, or in some cases even replace systematic sampling so that total sampling resources can be allocated over longer timeframes and larger spatial extents (Figure 3). In this way the combination of both systematic and opportunistic sampling can collect more data on abundance, distribution, and population demographics over time for use in IPMs.

Citizen science datasets are valuable for improving our understanding and monitoring of animal populations towards the goal of successful management and conservation (McKinley et al., 2017). The next priority should be the development of statistical models including IPMs that integrate opportunistic datasets. Here, we have demonstrated that citizen science projects, such as iSeeMammals, have the

ability to collect opportunistic datasets that cover large spatial regions and timeframes that can be used to improve the precision and accuracy of estimates of population parameters and vital rates.

## Acknowledgements

Many thanks to the iSeeMammals citizen scientists and dedicated staff at the New York State Department of Environmental Conservation who have assisted with data collection.

## Author contributions

CS, AF, and JH conceived the idea; CS and AF designed and implemented the citizen science project; CS analyzed the data; CS led the writing of the manuscript. All authors contributed critically to the drafts and gave final approval for publication. Any use of trade, firm, or product names is for descriptive purposes only and does not imply endorsement by the U.S. Government.

## References

Ashcroft, M.B., Chisholm, L.A., French, K.O., 2009. Climate change at the landscape scale: predicting fine-grained spatial heterogeneity in warming and potential refugia for vegetation. Glob. Change Biol. 15, 656–667. https://doi.Org/10.llll/j.1365-2486.2008.01762.x

Bain, K., Wayne, A., Bencini, R., 2015. Risks in extrapolating habitat preferences over the geographical range of threatened taxa: a case study of the quokka (Setonix brachyurus) in the southern forests of Western Australia. Wildl. Res. 42, 334–342. https://doi.org/10.1071/WR14247

Beirne, C., Lambin, X., 2013. Understanding the determinants of volunteer retention through capture-recapture analysis: answering social science questions using a wildlife ecology toolkit. Conserv. Lett. 6, 391–401. https://doi.org/10.llll/conl.12023

Beissinger, S.R., Westphal, M.I., 1998. On the use of demographic models of population viability in endangered species management. J. Wildl. Manag. 62, 821–841. https://doi.org/10.2307/3802534

Belant, J.L., Stappen, J.F.V., Paetkau, D., 2005. American black bear population size and genetic diversity at Apostle Islands National Lakeshore. Ursus 16, 85–92. https://doi.org/10.2307/3873061

Bonney, R., Cooper, C.B., Dickinson, J., Kelling, S., Phillips, T., Rosenberg, K.V., Shirk, J., 2009. Citizen science: a developing tool for expanding science knowledge and scientific literacy. BioScience 59, 977–984. https://doi.Org/10.1525/bio.2009.59.ll.9

Bowser, A., Hansen, D., He, Y., Boston, C., Reid, M., Gunnell, L., Preece, J., 2013. Using Gamification to inspire new citizen science volunteers, in: Proceedings of the First International Conference on Gameful Design, Research, and Applications, Gamification ’13. ACM, New York, NY, USA, pp. 18–25. https://doi.org/10.1145/2583008.2583011

Chandler, R.B., Clark, J.D., 2014. Spatially explicit integrated population models. Methods Ecol. Evol. https://doi.org/10.llll/2041-210X.12153

Dickinson, J.L., Shirk, J., Bonter, D., Bonney, R., Crain, R.L., Martin, J., Phillips, T., Purcell, K., 2012. The current state of citizen science as a tool for ecological research and public engagement. Front. Ecol. Environ. 10, 291–297. https://doi.org/10.1890/110236

Dickinson, J.L., Zuckerberg, B., Bonter, D.N., 2010. Citizen science as an ecological research tool: challenges and benefits. Annu. Rev. Ecol. Evol. Syst. 41, 149–172. https://doi.org/10.1146/annurev-ecolsys-102209-144636

Drewry, J.M., Van Manen, F.T., Ruth, D.M., 2013. Density and genetic structure of black bears in coastal South Carolina. J. Wildl. Manag. 77, 153–164. https://doi.org/10.1002/jwmg.443

Eveleigh, A., Jennett, C., Blandford, A., Brohan, P., Cox, A.L., 2014. Designing for dabblers and deterring drop-outs in citizen science, in: Proceedings of the SIGCHI Conference on Human Factors in Computing Systems, CHI ’14. ACM, New York, NY, USA, pp. 2985–2994. https://doi.org/10.1145/2556288.2557262

Follett, R., Strezov, V., 2015. An analysis of citizen science based research: usage and publication patterns. PloS One 10, e0143687. https://doi.org/10.1371/journal.pone.0143687

lacovides, I., Jennett, C., Cornish-Trestrail, C., Cox, A.L., 2013. Do games attract or sustain engagement in citizen science?: a study of volunteer motivations, in: CHI ’13 Extended Abstracts on Human Factors in Computing Systems, CHI EA’13. ACM, New York, NY, USA, pp. 1101–1106. https://doi.org/10.1145/2468356.2468553

Kéry, M., Gardner, B., Monnerat, C., 2010. Predicting species distributions from checklist data using site-occupancy models. J. Biogeogr. 37, 1851–1862. https://doi.Org/10.llll/j.1365-2699.2010.02345.x

McKinley, D.C., Miller-Rushing, A.J., Ballard, H.L., Bonney, R., Brown, H., Cook-Patton, S.C., Evans, D.M., French, R.A., Parrish, J.K., Phillips, T.B., Ryan, S.F., Shanley, L.A., Shirk, J.L., Stepenuck, K.F., Weltzin, J.F., Wiggins, A., Boyle, O.D., Briggs, R.D., Chapin, S.F., Hewitt, D.A., Preuss, P.W., Soukup, M.A., 2017. Citizen science can improve conservation science, natural resource management, and environmental protection. Biol. Conserv., The role of citizen science in biological conservation 208, 15–28. https://doi.Org/10.1016/j.biocon.2016.05.015

Nagy, C., Bardwell, K., Rockwell, R.F., Christie, R., Weckel, M., 2012. Validation of a citizen science-based model of site occupancy for eastern screech owls with systematic data in suburban New York and Connecticut. Northeast. Nat. 19,143–158. https://doi.org/10.1656/045.019.s611

Nichols, J.D., Williams, B.K., 2006. Monitoring for conservation. Trends Ecol. Evol. 21, 668–673. https://doi.Org/10.1016/j.tree.2006.08.007

Nov, O., Arazy, O., Anderson, D., 2014. Scientists@Home: what drives the quantity and quality of online citizen science participation? PLoS ONE 9, e90375. https://doi.org/10.1371/journal.pone.0090375

Pocock, M.J.O., Tweddle, J.C., Savage, J., Robinson, L.D., Roy, H.E., 2017. The diversity and evolution of ecological and environmental citizen science. PLOS ONE 12, e0172579. https://doi.org/10.1371/journal.pone.0172579

Pollock, K.H., 1982. A capture-recapture design robust to unequal probability of capture. J. Wildl. Manag. 46, 752–757. https://doi.org/10.2307/3808568

Robinson, O.J., Ruiz-Gutierrez, V., Fink, D., Meese, R.J., Holyoak, M., Cooch, E.G., 2018. Using citizen science data in integrated population models to inform conservation decision-making. bioRxiv 293464. https://doi.org/10.1101/293464

Royle, J.A., Young, K.V., 2008. A hierarchical model for spatial capture-recapture data. Ecology 89, 2281–2289. https://doi.org/10.2307/27650753

Sun, C.C., Fuller, A.K., Hare, M.P., Hurst, J.E., 2017. Evaluating population expansion of black bears using spatial capture-recapture. J. Wildl. Manag. 81, 814–823. https://doi.org/10.1002/jwmg.21248

Tenan, S., Pedrini, P., Bragalanti, N., Groff, C., Sutherland, C., 2017. Data integration for inference about spatial processes: A model-based approach to test and account for data inconsistency. PLOS ONE 12, e0185588. https://doi.org/10.1371/journal.pone.0185588

Williams, B.K., Nichols, J.D., Conroy, M.J., 2002. Analysis and management of animal populations, 1 edition, ed. Academic Press, San Diego.

Wolf, C., Ripple, W.J., 2017. Range contractions of the world’s large carnivores. R. Soc. Open Sci. 4, 170052. https://doi.org/10.1098/rsos.170052

Wood, C., Sullivan, B., Iliff, M., Fink, D., Kelling, S., 2011. eBird: Engaging Birders in science and conservation. PLoS Biol 9, el001220. https://doi.org/10.1371/journal.pbio.1001220

Xue, Y., Davies, I., Fink, D., Wood, C., Gomes, C.P., 2016. Avicaching: A Two stage game for bias reduction in citizen science, in: Proceedings of the 2016 International Conference on Autonomous Agents & Multiagent Systems, AAMAS ’16. International Foundation for Autonomous Agents and Multiagent Systems, Richland, SC, pp. 776–785.

Yackulic, C.B., Chandler, R., Zipkin, E.F., Royle, J.A., Nichols, J.D., Campbell Grant, E.H., Veran, S., 2013. Presence-only modelling using MAXENT: when can we trust the inferences? Methods Ecol. Evol. 4, 236–243. https://doi.org/10.llll/2041-210×.12004

Zipkin, E.F., Saunders, S.P., 2018. Synthesizing multiple data types for biological conservation using integrated population models. Biol. Conserv. 217, 240–250. https://doi.Org/10.1016/j.biocon.2017.10.017

